# Modeling long lifespans in eusocial insect populations

**DOI:** 10.1101/408211

**Authors:** Donepudi RaviTeja, Ramakrishna Ramaswamy

**Affiliations:** Department of Biotechnology and Bioinformatics, University of Hyderabad, Hyderabad, India; School of Computational and Integrative Sciences, Jawaharlal Nehru University, New Delhi, India

**Author notes:** Corresponding author; (RR).

## Abstract

Along with division of labour, and life-history complexities, a characteristic of eusocial insect societies is the greatly extended lifespan for queens. The colony structure reduces the extrinsic mortality of the queen, and according to classical evolutionary theories of ageing, this greatly increases the lifespan. We explore the relationship between the evolution of longevity and the evolution of eusociality by introducing age-structure into a previously proposed evolutionary model and also define an associated agent-based model. A set of three population structures are defined: (i) solitary with all reproductive individuals, (ii) monogynous eusocial with a single queen, and (iii) polygynous eusocial, with multiple queens.

In order to compare the relative fitnesses we compete all possible pairs of these strategies as well as all three together, analysing the effects of parameters such as the probability of progeny migration, group benefits, and extrinsic mortality on the evolution of long lifespans. Simulations suggest that long lifespans appear to evolve only in eusocial populations, and further, that long lifespans enlarge the region of parameter space where eusociality evolves. When all three population strategies compete, the agent-based simulations indicate that solitary strategies are largely confined to shorter lifespans. For long lifespan strategies the solitary behaviour results only for extreme (very low or very high) migration probability. For median and small values of migration probability, the polygynous eusocial and monogynous eusocial strategies give advantage to the population respectively. For a given migration probability, with an increase in lifespan, the dominant strategy changes from solitary to polygynous to monogynous eusociality. The evolution of a long lifespan is thus closely linked to the evolution of eusociality, and our results are in accord with the observation that the breeding female in monogynous eusocial species has a longer lifespan than those in solitary or polygynous eusocial species.

## Introduction

For living organisms death results from both extrinsic causes such as predation, disease, or accident, and intrinsic causes that include senescence. The force of natural selection decreases with age, with genes that affect later life being under reduced selection pressure [1–11]. As a consequence, according to mutation accumulation theory, alleles with deleterious effects — mutations — accumulate at later ages leading to senescence [2–7]. An alternate view put forth in the theory of antagonistic pleiotropy [4, 7, 12] is that senescence evolves due to side-effects that are deleterious when occurring late in life history, but which are favourable early on in the life cycle. These hypotheses propose that ageing can be evolved [11, 13, 14] and further, that an increase (or decrease) in extrinsic death-rate will increase (or decrease) the rate of ageing [1, 7, 15]. There are, of course, additional and more complex effects that are density and life-history dependent [1]. Some recent empirical and experimental observations give results contrary to simple classical predictions [16, 17] and the causes are largely unknown [18]. To explain the more complex effects and deviations, newer modeling approaches such as simulated annealing optimization [18], age-structured evolutionary models [19], hierarchical models [20] are being employed.

Eusociality, an advanced form of social organisation seen in several insect societies, is defined by reproducing (queen) and non-reproducing (worker) individuals that cooperatively care for the young [21–23]. Reproductive strategies depend on whether a colony can contain one or more queens: in monogynous species, each queen initiates a new and independent colony, while in polygynous species several queens remain in an established colony [8, 24–26]. Further, queens in monogynous eusocial societies have extraordinarily long lifespans as compared to those in polygynous species or compared to individuals of solitary species [8, 26–28]. The differences are striking: queens of the polygynous *Formica polyctena* and the monogynous *Formica exsecta*, both mound-building wood ants live 5 and 20 years respectively, while individuals of solitary species live at most for a month [8].

The interrelation between these two features has been studied previously [8, 20, 29] and indeed the occurrence of long lifespans in eusocial queens that also have a lower extrinsic death-rate is seen as a test of evolutionary theories [8]. Shorter lifespans that are associated with polygyny are understood within the framework of evolutionary theory [8] under the assumption that polygynous species have a higher mortality risk of queens since they inhabit fragile nests that increase the extrinsic mortality. However, there are polygynous species that inhibit permanent nests with the lifespan of those queens being lower than a related monogynous queen [10], suggesting that the link between queen lifespan and eusociality needs further exploration. In this context to explore the interdependency in evolution of these features, newer evolutionary models are required. These models should allow for various competing strategies of both eusociality and ageing.

The Nowak, Tarnita, and Wilson [30] (NTW) model for the evolution of eusociality examines the population structure of eusocial organisms in order to understand the conditions for evolution of eusociality. The details of the model are as follows: The eusocial population consists of several colonies, each of which consists of a reproductive *queen* and non-reproductive workers. The total number of queens and workers in a colony is its size. Colonies of a particular size show similar behaviour (namely the death- and birth-rates). The queen and workers of a colony die at a characteristic death-rate while queens reproduce at a characteristic birth-rate (these rates are size dependent). Progeny either stay in the nest with probability *q* to become workers and increase the size of the parent colony by one, or migrate with probability (1-*q*) to start a new colony. Death of the queen kills the entire colony while death of a worker reduces the colony size by one (Fig 1.c). The model assumes that the benefits of a colony are realised only after the size of the colony reaches a threshold above which it stays constant. Solitary populations, on the other hand, consist of homogenous individuals which can reproduce and die at characteristic rates (Fig 1a). NTW compete the two populations and examine conditions when natural selection favours the eusocial allele. When large benefits are associated with colonies above a particular size, eusociality can evolve for a range of *q* by increasing the rate of oviposition and reducing the death-rate for queens. Ageing is absent in the model, the rate of oviposition and/or death-rate being age-invariant. There has been some debate on the conclusions that can be drawn from the initial studies [31].

**Figure 1.**
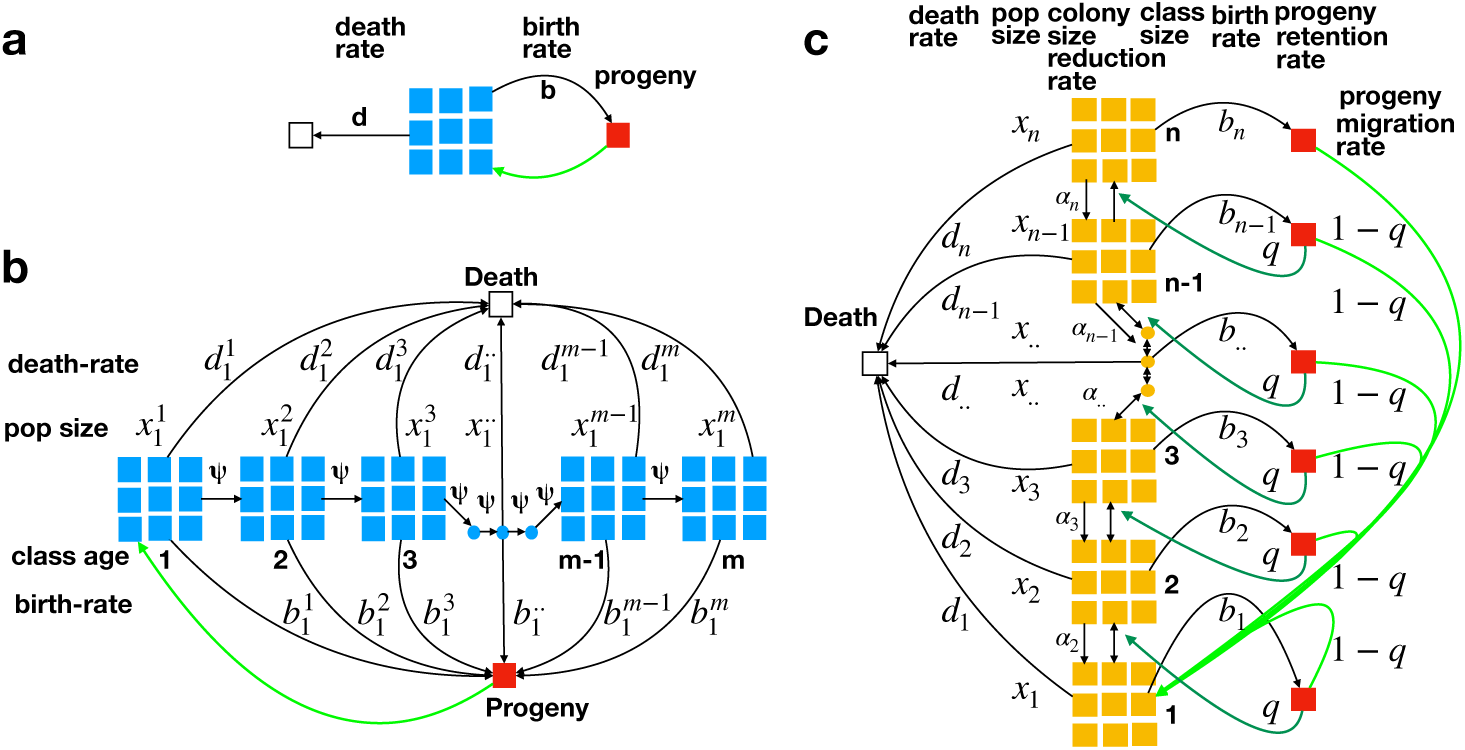
Graphical depictions of various evolutionary models. A single reproductive individual is represented by a blue square, the eusocial colony by an orange square, and red square represent progeny. Subscripts on variables denotes the size of colony, and superscripts denote the age of the queen. a) Solitary model: individuals reproduce at rate *b*, die at rate *d*, and the progeny joins the population. b) Age structured solitary model: the populations is segregated into age-classes, each of which move to next age-class at rate *Ψ*. All individuals reproduce at rate 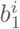 and die at rate 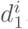, and the progeny joins the first age class. c) Monogynous eusocial model (NTW): The eusocial colonies are segregated into size-classes and the queen of each colony reproduces at rate 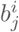 and die at rate 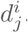. The progeny can remain to increase the size of the parent colony by 1 or start a new colony of size 1. Workers in a colony reduce the size by 1 upon death.

Does the eusocial population structure leads to the evolution of long lifespans in queens? Keeping this underlying question in mind, the aim of the present work is to test the conditions for evolution of long lifespans in such eusocial populations. The NTW model operates within the standard theory of natural selection, making it possible to evaluate multiple competing hypotheses [30, 31]. We extended the NTW framework to build two models, one which introduces age-dependence in the reproductivity and one that uses agent based modeling to allow for *polygynous eusocial* life-history, in addition to solitary and monogynous eusociality. We use both models to determine how the population structure, and specific parameters affect the evolution of long lifespans, and how this can in turn affect the evolution of eusociality.

## Model

We first briefly review the essential components of the models studied by NTW. For a solitary population model the abundance of solitary individuals, denoted *x*, evolves in time according to the dynamics

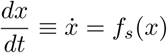

where the *t* is time and the function *f*_*s*_ contains details of the assumed models of birth and death. In the simplest case of constant birth-rate *b* and death-rate *d, f*_*s*_(*x*) = (*b - d*)*x* (Fig 1a).

In an eusocial population model, one considers *x*_*j*_ colonies (or nests) of size *j* with *j* = 1,2,*…, n*. Where *n* is the largest nest size. The total population is

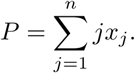

The population of a nest changes by growth and by migration, namely

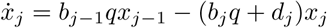

with *j* = 2, 3, *…*, while for nests of size 1 the dynamics is

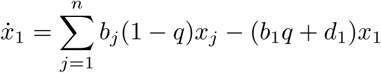

where *b*_*b*_ and *d*_*d*_ are the birth and death rates for colonies of size j. The probability that progeny migrate from the parent colony is 1–*q*; see Fig 1.c.

Features such as density limitations and worker mortality can be included in the model and NTW have extensively studied these. Our present interest is in the inclusion of age-dependence within this general framework. We discuss this variation next, first within the NTW model, and thereafter in an agent-based model that has a greater level of flexibility in implementing the population structure. Parameters and variables used in these models are summarised in Table 1.

**Table 1.** Summary of the model variables and parameters.

| Variable | Description | Comment |
| --- | --- | --- |
| $x$ | Population size | Total population for solitary NTW model. |
| $b$ | Birth-rate | Rate of progeny birth for solitary NTW model. |
| $d$ | Death-rate | Rate of death in solitary NTW model. |
| $n$ | Largest nest size | Size of the largest nest in eusocial models. Chosen as 30. |
| $x_j$ | Size of size-class | Number of colonies of size $j$ in eusocial NTW model. |
| $P$ | Population size | Total population for eusocial NTW model. |
| $b_j$ | Birth-rate | Rate of progeny birth for colony of size $j$ in an eusocial NTW model. |
| $d_j$ | Death-rate | Rate of death of queens for colony of size $j$ in an eusocial NTW model. |
| $q$ | Stay put probability of progeny | In eusocial populations, it is the rate of progeny to stay in the parent colony to become a worker. |
| $\beta$ | Effective birth-rate | Rate of progeny birth for a continuous model (effective birth-rate when the intrinsic death-rate is subtracted). |
| $\mu$ | Extrinsic Death-rate | Rate of extrinsic death in the continuous age-structured population model. |
| $S$ | Population size | Total population at a particular time for continuous age-structured model. |
| $m$ | Maximum age | It is the last age in finite age population models, chosen as 30. |
| $r$ | Maximum reproductive age | The age or age-group after which birth-rate is equal to zero. |
| $K$ | Lifetime reproductive capacity | Total number of progeny produced in its lifetime. Assuming lifetime investment in reproduction is same across comparing models, it is considered a constant. |
| $h_1$ | Birth-rate at first age | The starting age or age-group birth-rate. |
| $h_2$ | Birth-rate at age group $r$ | In finite age population models, this is the birth-rate at last reproductive age or age-group $r$. |
| $b_j^i$ | Discrete model Birth-rate | Rate of progeny birth (effective birth-rate when intrinsic death-rate is subtracted) for age-structured NTW model. Depends on age ($i$) and size ($j$) of the colony. |
| $d_j^i$ | Extrinsic death-rate | Rate of extrinsic death of queens in age-structured NTW model. Low enough to avoid extinction. Depends on age ($i$) and size ($j$) of the colony. |
| $g$ | Group (colony) benefit | Defines the benefits due to colony structure. In our simulations this takes values in the interval $[1,10]$. |
| $x_j^i$ | Size of an age-size class | Number of nests in particular age ($i$)-size ($j$) class. It denotes the number of colonies of size $j$ and with queen of age $i$. |
| $\psi$ | Time progression rate | Rate of transfer of colonies to the next age class. Chosen as 0.25, does-not affect results qualitatively. |
| $\alpha_j$ | Colony size reduction rate | Product of worker death rate and number of workers in the colony. It defines the rate at which colonies move to a lower size-class. |
| $\phi$ | Density dependence factor | In age-structured NTW models, it is defined as $1/(1 + U)$, where $U$ is the total population and can be equal to $x_E$ or $x_S$ or $x_E + x_S$. |
| $x_S$ | Population size (fitness) | Total population of age-structured solitary NTW model and is the absolute fitness of that population. |
| $x_E$ | Population size (fitness) | Total population of age-structured eusocial NTW model and is the absolute fitness of that population. |
| $p$ | Age at maximum birth-rate | In a unimodal (tent-shaped) reproductive strategy, the birth-rate has a maximum at an intermediate age. |

### Age-dependence in evolutionary dynamics models

To understand the effect of age-structure on a solitary population we define a continuous population model [32]. The results can then be compared with the age-structured NTW model. It is important to include the effect of age *a* in order to incorporate different breeding patterns, and we first obtain an expression that defines the fitness of a population in terms of the ageing strategy. For a population with finite age structure, the critical threshold *S* (or fitness) for growth is defined as [32]

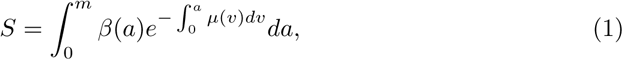

where *β*(*a*) is an age-dependent effective birth-rate, *m* is the maximum age and the rate of death due to extrinsic causes is *µ*. Assuming that this is an age-independent constant *D* gives

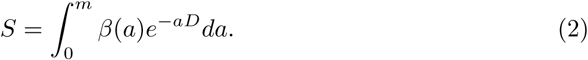

The functional dependence of the effective birth-rate, namely *β*(*a*) defines the “ageing strategy” of the population. We consider *β*(*a*) to be linear, interpolating from an initial rate of *β*(0) = *h*_1_ to a final *β*(*r*) = *h*_2_ with *β*(*a*) = 0 beyond, namely for *a > r* (Fig. 2a). We chose this function and later a tent function (with an additional parameter *p*) to define ageing so to approximate the unimodal ageing behaviour seen in nature, where the effective birth-rate starts at *β*(0) = *h*_1_ first increases till age *p* and then decreases by age *r* (*β*(*r*) = *h*_2_) (Fig. 2b). Choices of *h*_1_, *h*_2_ and *r* will specify the ageing strategy (Fig. 2a) in the following way. For any form of *β*(*a*), the total reproductivity is kept constant at *K* [33]. Thus

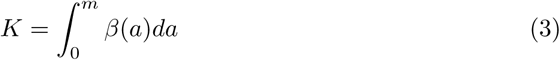

**Figure 2.**
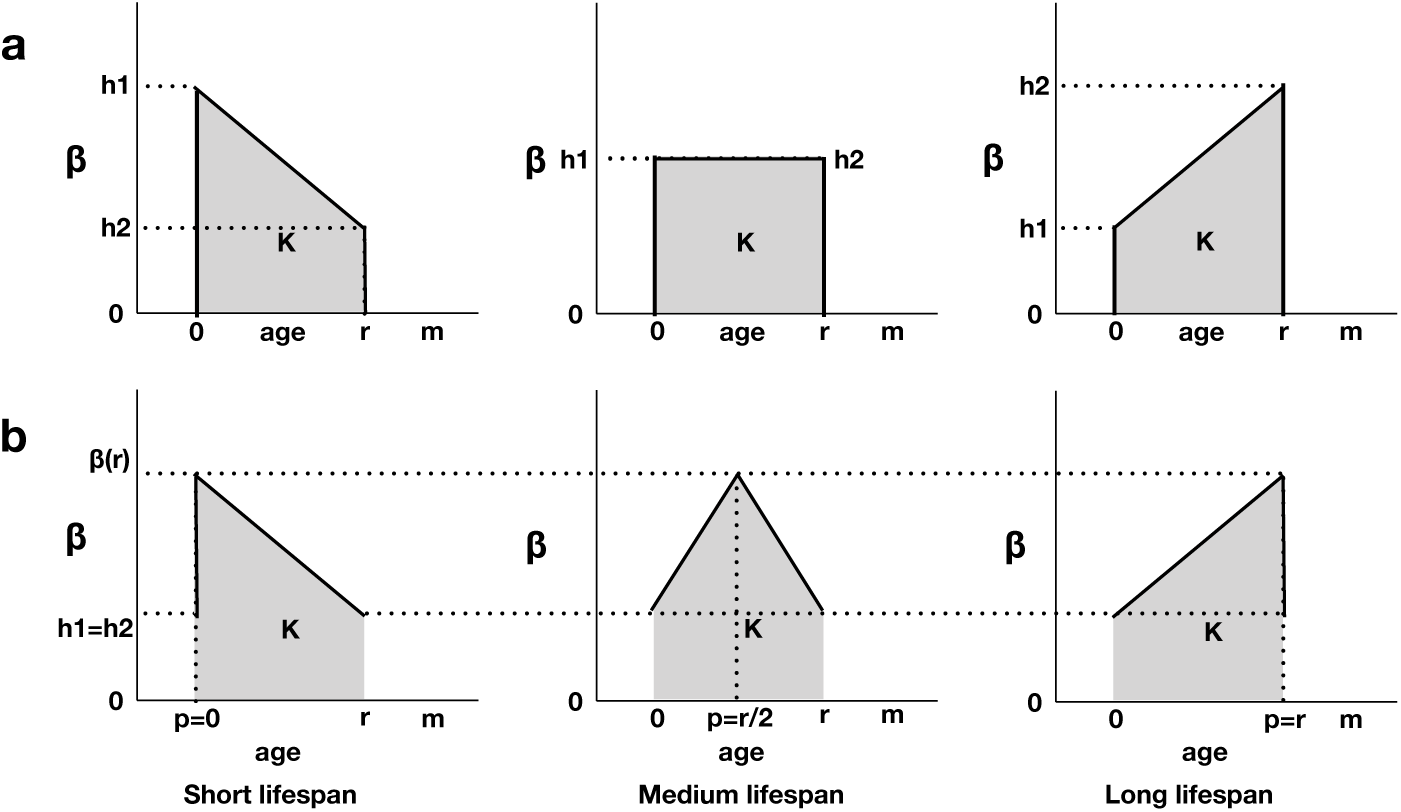
Graphical depictions of ageing-strategies. *β* denotes the effective birth-rate a) Simple monotonous ageing strategy and b) a unimodal (tent shaped) ageing strategy.

For the case above, namely *β*(*a*) = 0 for *a > r* and linear in [0, *r*] with *β*(0) = *h*_1_, *β*(*r*) = *h*_2_, we get

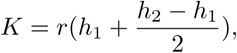

which gives

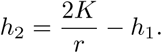

Thus for fixed *K* and *r*, either *h*_1_ or *h*_2_ can be chosen. The optimal strategy, namely the maximal *S* for a constant life-time reproductive capacity *K* defines a variational problem (within the linear choice for *β*). Solving Eq. (2) using this condition yields

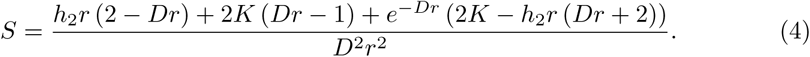

For given *K, r* and *D, S* will reduce with increasing *h*_2_: the optimal strategy is to have higher early effective birth-rate. A population of solitary individuals thus cannot achieve a long-life life history. As effective birth-rate is defined as the difference between birth-rate and intrinsic death-rate, the ageing strategy can be achieved by various combinations of birth-rate and death-rate functions.

We include ageing structure within the NTW model for the evolutionary dynamics of solitary and eusocial organisms [30] as follows.

- The age structured NTW solitary population is segregated into classes identified by age and size. Here 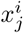 is the number of groups of age class *i* = 1, 2, *…* and size *j*, with *j*=1 being used to denote a solitary population. The ageing strategy of the population will define the birth 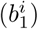 rates of each age class. Here and in the rest of this paper, the effective birth-rate 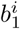 has had the intrinsic death-rate subtracted. The extrinsic death-rate is denoted as 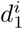. Individuals in an age class move to next age class (*i → i* + 1) at ageing rate *Ψ*, and progeny join the first age class (Fig 1b). For a population with *m* age classes, the master equation can be written as

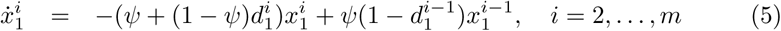

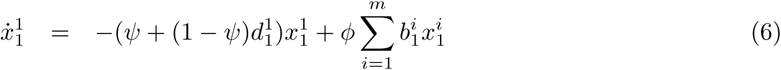

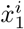 being the change in population for each age class other than the first. The starting age class is defined by 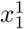 and 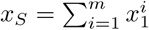 is the total population size.

- For an age structured NTW monogynous eusocial population, *j* ≥ 1. There are both queens and workers, with nests being segregated into age-size classes. The properties of each eusocial colony (birth-rate, death-rate etc.,) are dependent on its age-size class. The population size of each age-size class (number of colonies) is denoted as 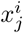, where age is given by *i* and colony size (number of workers and queen) is given by *j*. A monogynous eusocial colony has following six behaviours, queen dies at specific extrinsic death-rate 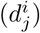 and the colony dies, 2) worker bee dies and colony size reduces by 1, 3) queen reproduces at birth-rate 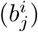, 4) the progeny stay in the colony at *q* and increases the colony size by 1, 5) the progeny migrate at 1–*q* to start a new colony of size 1 and not changing the size of the parent colony, 6) the queen age at ageing-rate (*Ψ*) and moves the colony to next age-class (*i → i* + 1). The combination of these behaviours define the dynamics of the population (Fig. 3).

**Figure 3.**
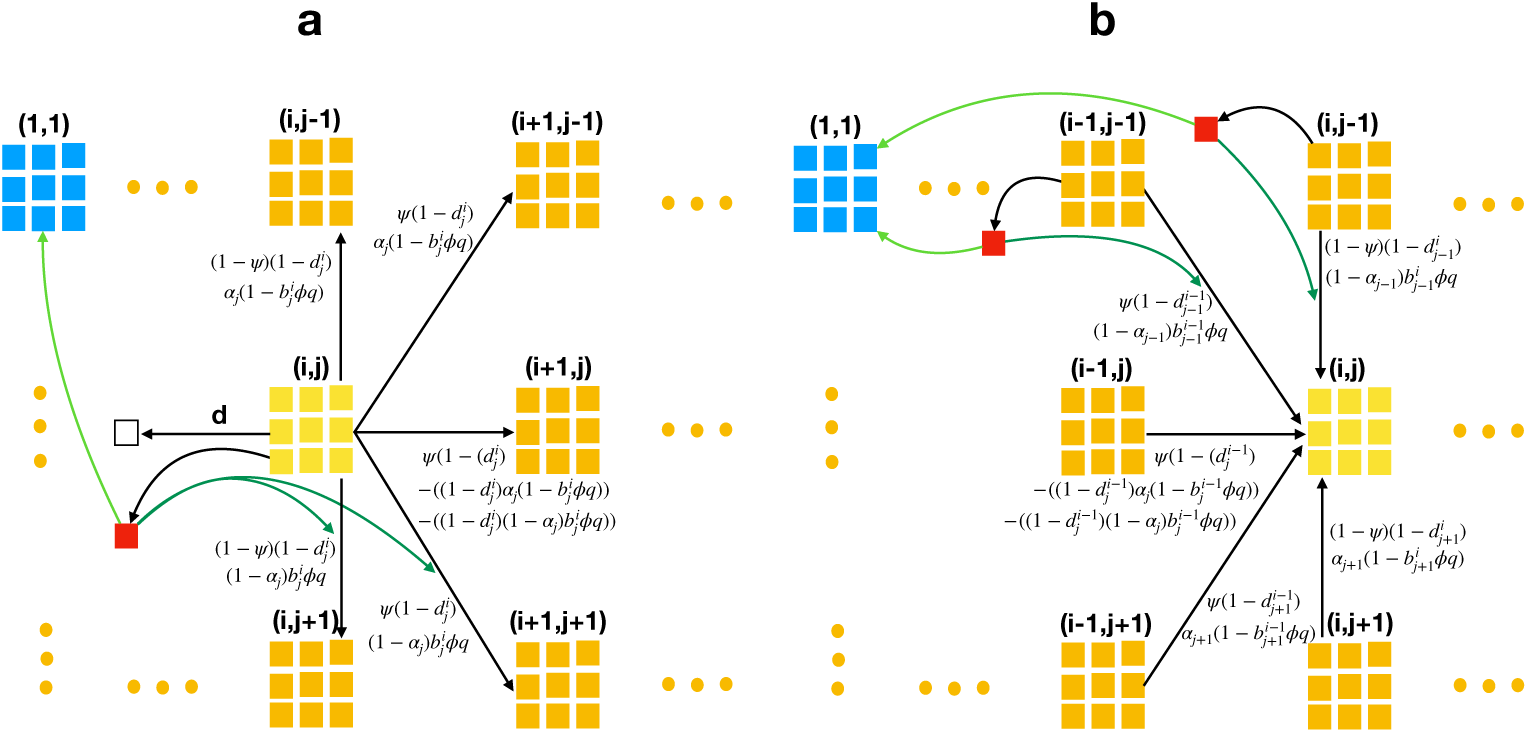
Graphical depiction of population dynamics in a monogynous eusocial model. A monogynous eusocial colony is indicated by an orange square. A group of yellow squares represents the age-size class whose dynamics are depicted. A blue square represents a colony of size 1, and a red squares represent progeny. The number of squares in a group is only for representation purposes and does not specify the size. The size of colony and the age class, namely the age of the queen is indicated above each group as an ordered pair (age,size). a) Depicts all possible dynamics for movement of colonies *out* of an age-size class, while b) illustrates all possible dynamics for movement of colonies *into* an age-size class.

The parameter *α*_*α*_, the product of individual worker death-rate and number of workers in the colony defines the rate at which one age-size class transitions to another class with smaller size. If *d*_*d*_ is the death-rate of workers in a colony of size *j*, then *α*_*j*_ = (*j–*1)*d*_*w,j*_. The appropriate master equation for a population of monogynous eusocial organisms is

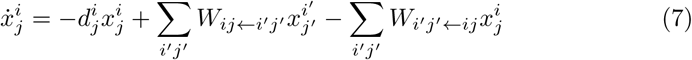

where the transition rates *W*_*ij←i*_*′* _*j*_*′* indicate the rate of movement from the age-size group *i*^*l*^*j*^*l*^ to the age-size group *ij*. Fig 3 for description of *W*_*ij←i*_*′* _*j*_*′* and *W*_*i*_*′* _*j*_*′*_*←ij*_:

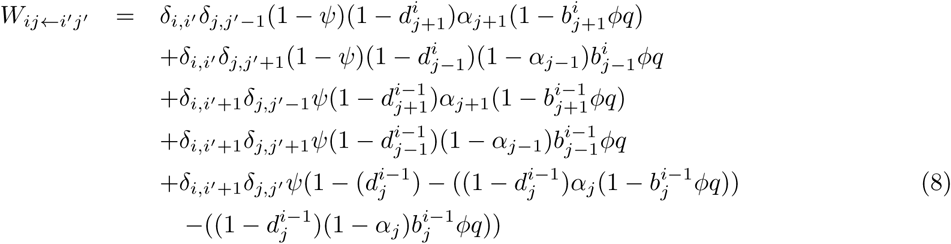

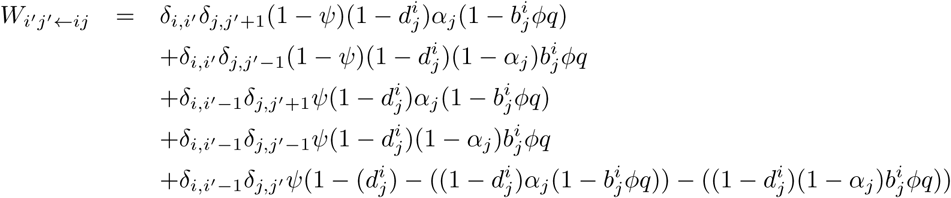

and *δ*_*δ*_ is the Kronecker delta namely 1 if *i* = *j* and 0 if *i ≠ j*. *ϕ* is the density limitation for growth of population, here taken to be *ϕ* = 1*/*(1 + *x*_*E*_) where *x*_*x*_ is the total population and is given by

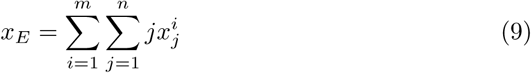

Where *m* is the largest age-group and *n* is the maximum colony size. We keep the extrinsic death-rate constant and birth-rate is a linear function of age (Fig 2a). For colonies with size above a threshold (*Th*), the birth-rate is multiplied by the group benefit *g*, and the death-rate by the factor 1*/g*.

Simulations were run separately for a population of eusocial organisms with a single queen (the monogynous case) as well as for a population of solitary organisms. In order to numerically solve the model, an age and colony size structured Leslie matrix *L* [34] of size *mn* × *mn* is constructed at each time-step *t* using the given transition rates. The population vector for for the eusocial model 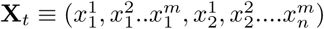 has *mn* elements. The population vector at time *t*+1 is given by

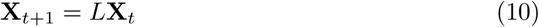

Given the population vector **X**, the population size (*x*_*E*_) at each time can be calculated using Eq. 9. For solitary models *m*=1 and the population measured is denoted *x*_*S*_. \The above model is numerically solved using a standard R [35] code for a range of *h*_2_ and *D* (the constant extrinsic death-rate) values until steady state is achieved. The code has been provided in the Supplementary Material.

In order to examine the effect of ageing on eusociality, we simulate both the monogynous eusocial model and the solitary models together, where the density limitation factor *ϕ* depends on the total population *x*_*E*_ + *x*_*S*_ and is given by *ϕ* = 1*/*(*x*_*E*_ + *x*_*S*_). Such competition experiments are conducted for a range of *h*_2_, *q* and *g* values. When either *x*_*E*_ or *x*_*S*_ falls below a specified level, that particular strategy, namely eusociality or solitary, is deemed extinct.

### Agent-based Models

A population can in principle implement a number of strategies: adopting a solitary lifestyle, eusociality with a single queen, or eusociality with several queens (polygyny). We need to model a more complex population structure, and agent-based modeling allows for an easier implementation of this complexity. We therefore study an agent-based model for age-structured evolutionary dynamics using RNETLOGO [36] and NETLOGO [37]. All codes are available in the Supplementary Material.

The agent-based description of the above evolutionary model is extended to include the polygynous population structure as follows. There are two types of agents in the model, queens and workers. Reproductive females or queens age at a given rate and have specified birth and death-rates. Offspring stay within the parent colony at a given rate. Migrating progeny can start a new colony or join another colony, and there is a specific rate at which the non-migrant offspring are successfully established as queens. Workers, on the other hand, have an ageing process with a specific death-rate. The agents interact according to the rules outlined below.

Solitary species consists only of queens which reproduce or die at a defined age-dependent rate. Monogynous and polygynous eusocial strategies result in populations that are organised in colonies with both queen and worker agents. All queens and workers are part of any one of the colonies. Each offspring of a queen can either remain in order to become workers or leave to start a new colony in monogynous populations or in polygynous populations, can also join any other colony as additional queens at a specified successful establishment rate. The reproductive rate of a queen is colony size, namely on the total number of workers and age dependent; in polygynous colonies, the reproductive rate is scaled in order to account for competition between the queens. As in previously defined models, both short and long-term strategies are employed. In the short strategy, the effective birth-rate *b* starts from a high value, linearly reducing to a minimum at age *r*, while for the long term strategy, *b* starts from a low value and increases linearly (Fig 3a) till age *r* and for *a > r b* = 0. To make the ageing strategies more generic, a tent-shaped function is considered. Here the *h*_1_ and *h*_2_ represents the birth-rate at initial and at age group *r* and *p* represents the age group at which the birth-rate is maximum. As the life time reproductive capacity (*K*) and number of reproductive age groups (*r*) is constant across finite age models, and given *h*_1_, *h*_2_ are also equal and constant, the value of the birth-rate *b*(*r*) at any chosen *p* will be the same. Here *p* will define the ageing strategy (Fig 2b): smaller *p* indicates the short lifespan strategy and larger *p* the long lifespan strategy.

## Results and Analysis

The models discussed above in the previous Section have a finite age-structure, with the birth-rate and intrinsic death-rate being linear functions of age. The ageing function is the difference between the age-weighted birth- and intrinsic death-rates, and is represented as effective birth-rate function. For constant life-time reproductive capacity *K*, the larger the age-weighted effective birth-rate, the longer is the lifespan of the organism (Fig 2). These ageing strategies thus can be competed and a particular dominant strategy can emerge. Note that we define the fitness of a population as its size.

### Solitary Populations

Eqs. (1-4) and (5-7) represent the age-structured population model for solitary organisms, with the fitness given in Eq. (4). In our simulations, we take *K* = *r*= 20 and vary *h*_2_. Fig 4 shows the resulting fitness as a function of *h*_2_ and the death rate, *D*. Since the age-weighted average effective birth-rate 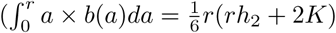 is proportional to *h*_2_, a larger value of *h*_2_ would correspond to the “long” ageing strategy. As expected, independent of the value of the extrinsic death-rate *D*, the fitness decreases monotonically with *h*_2_ showing that independent of the extrinsic death-rate, a short (semelparous) life history is inevitable for a simple population with the specified ageing function.

**Figure 4.**
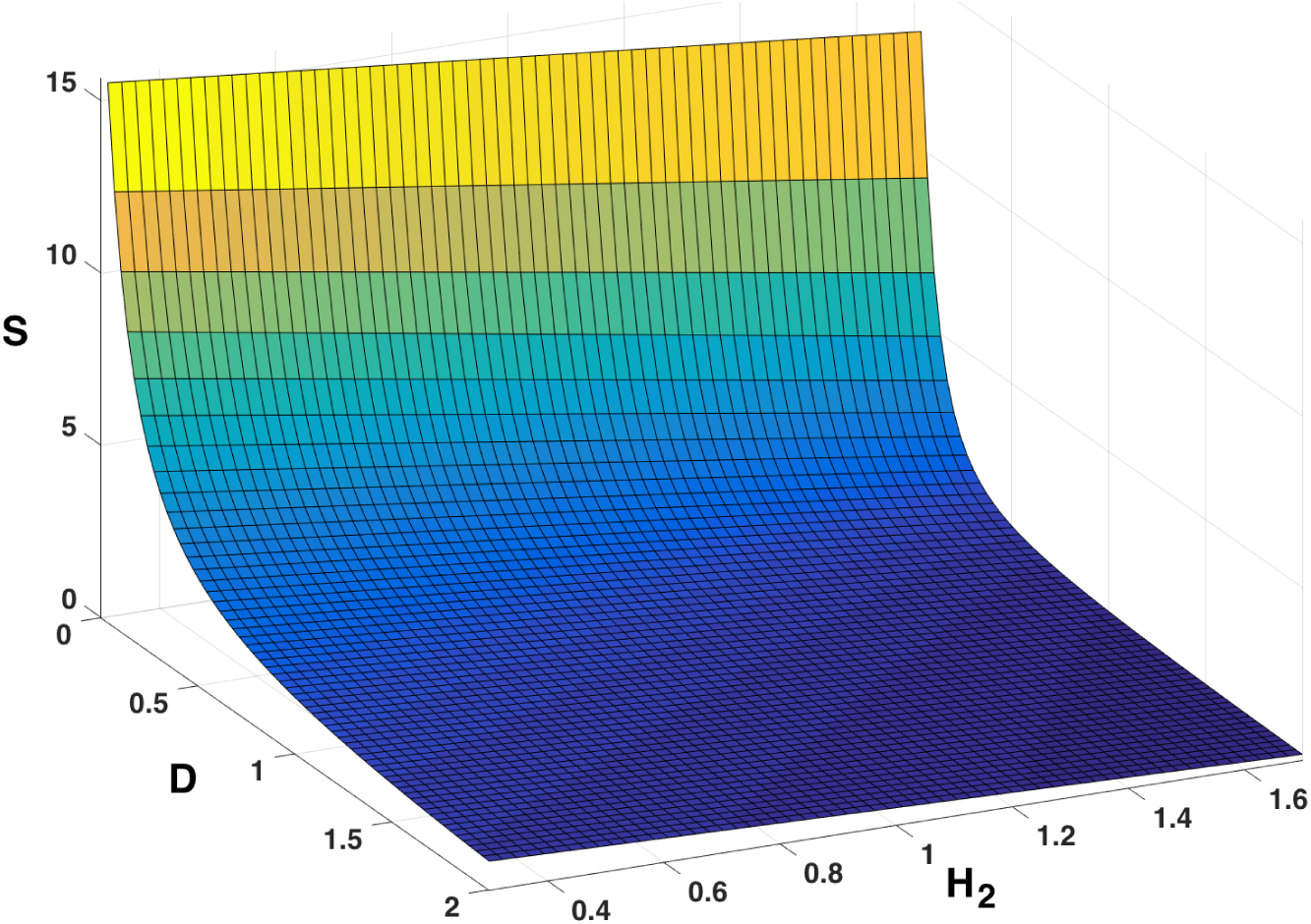
Fitness of the age-structured solitary population, plotted as a function of extrinsic death-rate *D* and *h*_2_ which decides the ageing strategy (see text).

A solitary population cannot reduce the ageing rate (namely increase the lifespan) due to reduced extrinsic death-rate under the conditions of these simulations. For solitary species, the short-term strategy always outcompetes a long term strategy, suggesting that long lifespans cannot be evolved in solitary species. We have also simulated Eq. (6): the asymptotic population size shows (Fig 5) that the short strategy *always* out-competes others. As above, we use a linear variation, with *h*_2_ defined in a similar manner: Similar results (not shown here) were achieved using agent-based model of solitary populations. This show the concurrence between our models.

**Figure 5.**
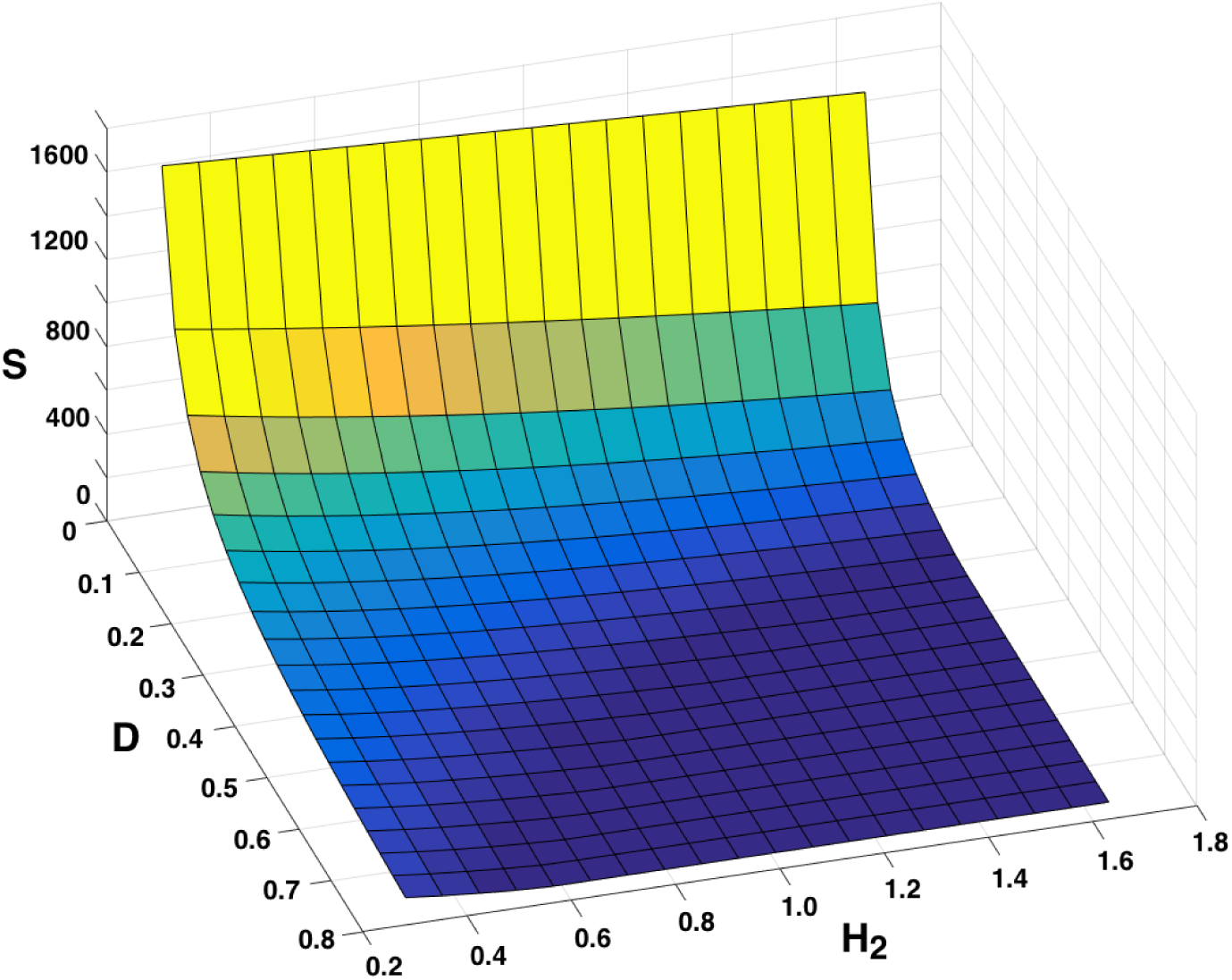
Fitness of the solitary age-structured evolutionary model, plotted as a function of the extrinsic deathrate *D* and *h*_2_ for the solitary evolutionary dynamics model. As in Fig 4a larger *h*_2_ corresponds to a long-life strategy.

### Eusocial Populations

Eq. (7-9) specifies the age-structured population model for monogynous eusocial organisms. As in the case of the solitary population, fitness is measured as the size of the population, namely Eq. (9) in steady state. The ageing function defined by *h*_2_ (Fig 2a) is used to normalise the birth-rate and/or death-rate, and a number of different strategies are modelled.

Fig 6 shows the fitness as a function of *h*_2_ and *D*, and here one can see a departure from the behaviour of solitary populations (Figs. 4 and 5). For low extrinsic death-rate, the “long” strategy has a higher fitness than the short strategy, and for high extrinsic death-rate, the opposite is true. Due to the characteristic shape of the fitness landscape, there are regions of non-monotonic fitness, namely for constant extrinsic death-rate the fitness is minimal at an intermediate value of *h*_2_, suggesting that fitness can grow both by increasing or decreasing the lifespan. It means that in a population started with a queens of specific life-span, the decrease or increase of extrinsic death-rate might not always lead to increase or decrease in life-span respectively and contraty can be possible.

**Figure 6.**
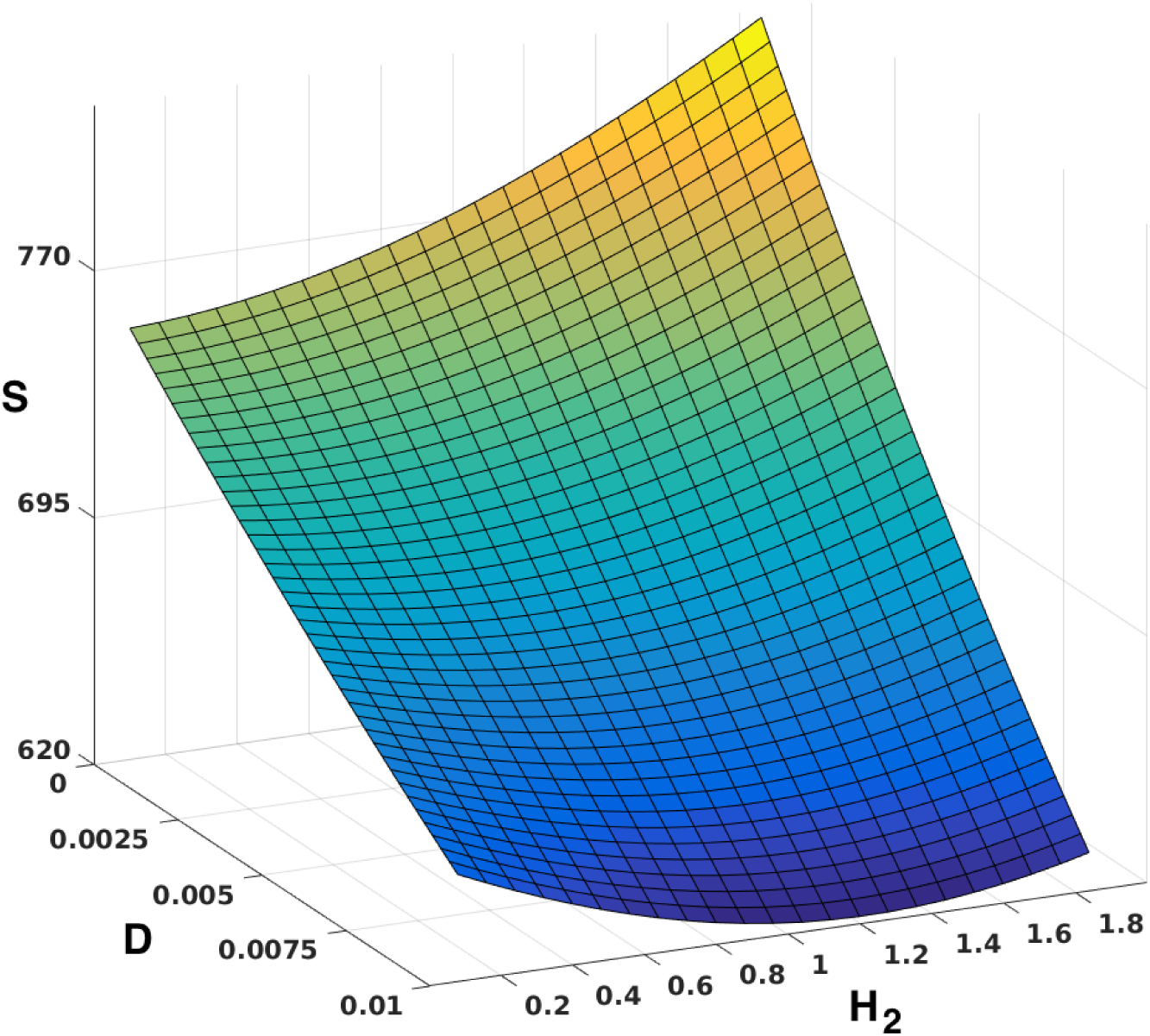
Fitness of the solitary age-structured evolutionary model. Fitness plotted as a function of extrinsic deathrate *D* and *h*_2_ for monogynous eusocial evolutionary dynamics model. *h*_2_ is proportional to the ageing strategy.

When we competed these monogynous eusocial strategies with one another (also using the agent-based model), we achieved results in accord with the above fitness landscape. When long and short monogynous strategies are competed the short strategy outcompetes the long when probability to stay *q* is small and the colony benefit *g* is large. On the other hand, the long strategy dominates for median and high values of *q* (data not shown). When polygynous eusocial strategies compete against each another in an agent-based model, we find that the short strategy dominates for a wider range of *q* than the monogynous populations. This suggests that long lifespans can also be evolved in polygynous eusocial populations although over a comparatively limited parameter range.

### Solitary vs Monogynous Eusocial strategies

We competed solitary strategy with the monogynous eusocial strategy using both evolutionary dynamics model and agent-based model. The simulations are carried out with an initially equal number of solitary and eusocial individuals in the population, and the dynamics are allowed to evolve for different *q, g* and *h*_2_ until one of the strategies dominate. Results are given in Fig 7: the dominant strategy is shown as a function of the three parameters. The monogynous eusocial strategy dominates over a larger range of *q* and *b* when long lifespans (larger *h*_2_) are considered as opposed to the case when shorter lifespan strategies (small *h*_2_) are considered.

**Figure 7.**
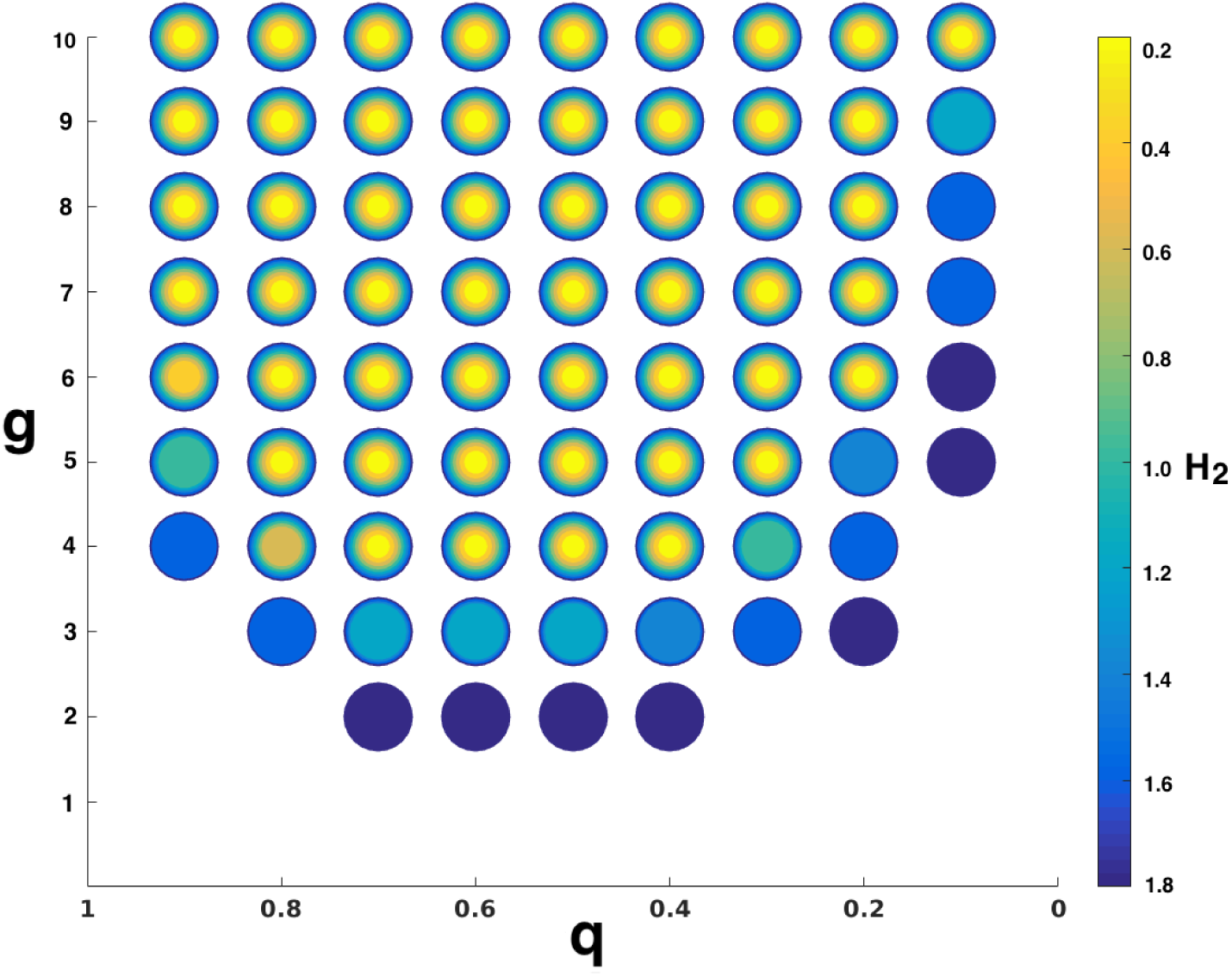
Competition between solitary and monogynous eusocial populations: Each filled circle denotes the evolution of monogynous eusociality. The colour of the filled circle denotes the value of *h*_2_. In competition, when long strategies are employed, monogynous eusociality evolves for lower *g* values than when short strategies are used.

The four strategies that are possible come from combinations of solitary versus monogynous eusocial populations with monotone decreasing birthrate (*h*_2_ *< h*_1_) or increasing (*h*_2_ *> h*_1_) namely the short or long ageing strategies (Fig 2a): these are denoted SS, ES, SL, and EL. Simulations are carried out for different *b* and *q* with equal numbers of all four types of individuals in the population initially. The system is allowed to evolve until a single strategy dominates: this is shown as a function of the two parameters in Fig 8. The solitary short (SS) and EL (eusocial-long) are the only strategies that eventually dominate, and the other two possibilities, namely SL, the late-breeding solitary populations or ES, eusocial populations which breed early are not seen in our simulations. For median ranges of the probability to stay *q*, we find that eusocial populations with a long period of queen fecundity dominates, while solitary populations with early reproduction are preferred when the probability to stay *q* is close to 0 or 1. The parameter range corresponding to the evolution of eusociality increases in comparison to the age-independent model studied in NTW [30]. Compare Figs 7, 8 with corresponding results in [30]. Within this model, therefore, this indicates that a long lifespan *promotes* eusociality: a monogynous eusocial population with a long-lived queen almost always outcompetes a similar society with a short-lived queen unless the probability to stay (*q*) is low and group benefits *g* are high and a solitary one unless the probability to stay (*q*) is at extremes.

**Figure 8.**
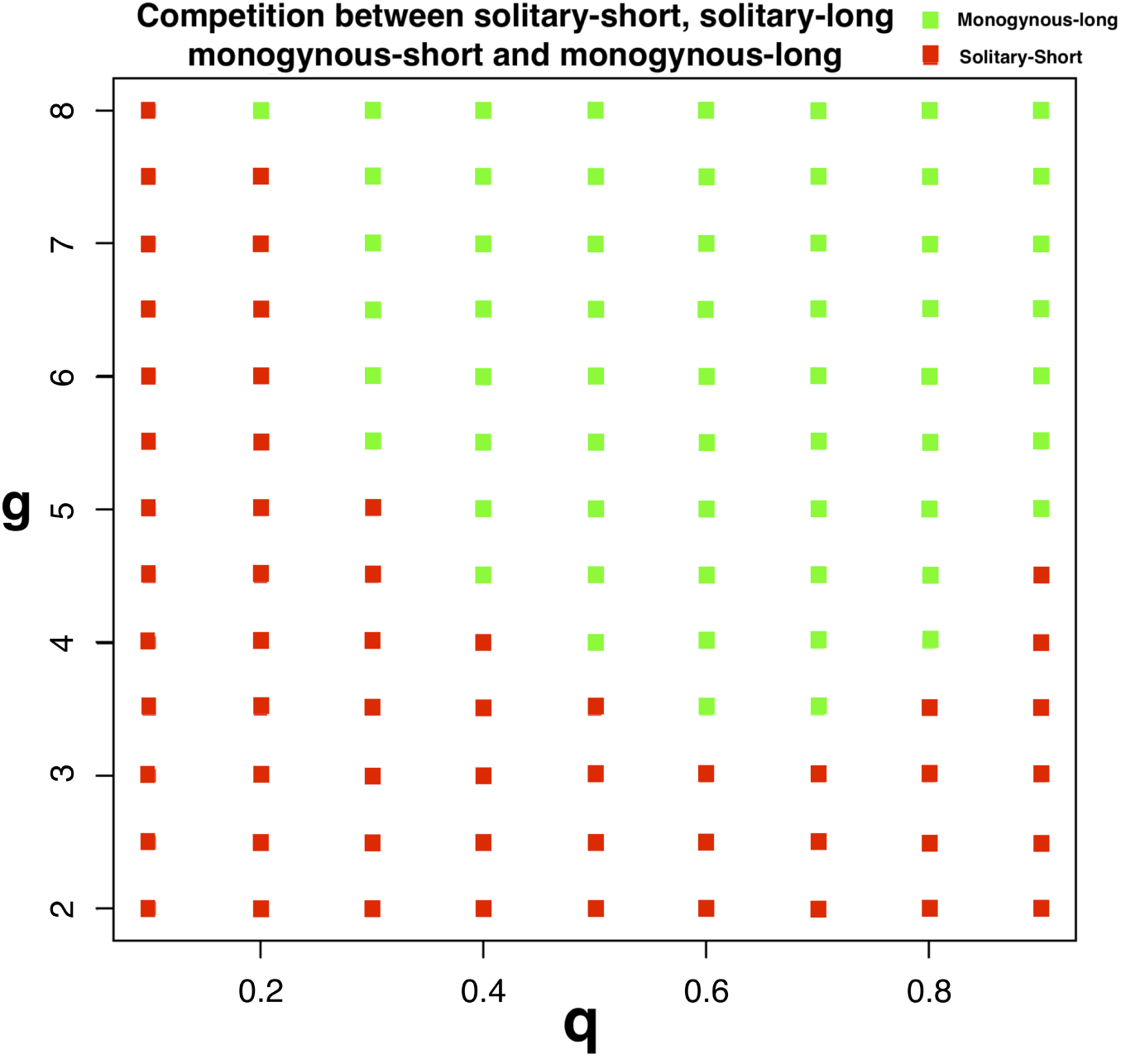
Competition between SS, SL, MS and ML strategies. Green boxes denote evolution of the monogynous long-lifespan (ML) and red boxes denote evolution of the solitary short-lifespan (SS) cases. As discussed in the text, SL and MS are not observed.

### Solitary vs Monogynous vs Polygynous eusocial strategies

If one allows the colony to be polygynous, there are two more strategies to be considered, namely PS and PL. Using the agent-based model, all six strategies are defined and simulations are carried out as described above; For chosen *b*, the SS strategy dominates for both low and high *q*, while the polygynous strategy PS dominates for intermediate *q*, and the monogynous is dominant at large *q* values (data not shown).

As discussed in previous section, one can consider more general age-fecundity structures (Fig 2b). If the rate of oviposition is taken to be maximal at reproductive age *p* (we took a piecewise linear function, increasing to a maximum at *p* with a subsequent linear decrease). Varying *q* and *p* but keeping the area under the curve constant we compete solitary and both eusocial strategies in order to determine the dominant strategy. As can be seen in Fig 9, the solitary strategy dominates for low values of *q* (decreasing somewhat with increasing *p*). For intermediate *q* the solitary strategy emerges only at low values of *p* while the polygynous strategy dominates for larger *p*. For higher probability to stay, as may be expected, the monogynous strategy dominates, except for the possibility for low *p* when the solitary strategy may be preferred.

**Figure 9.**
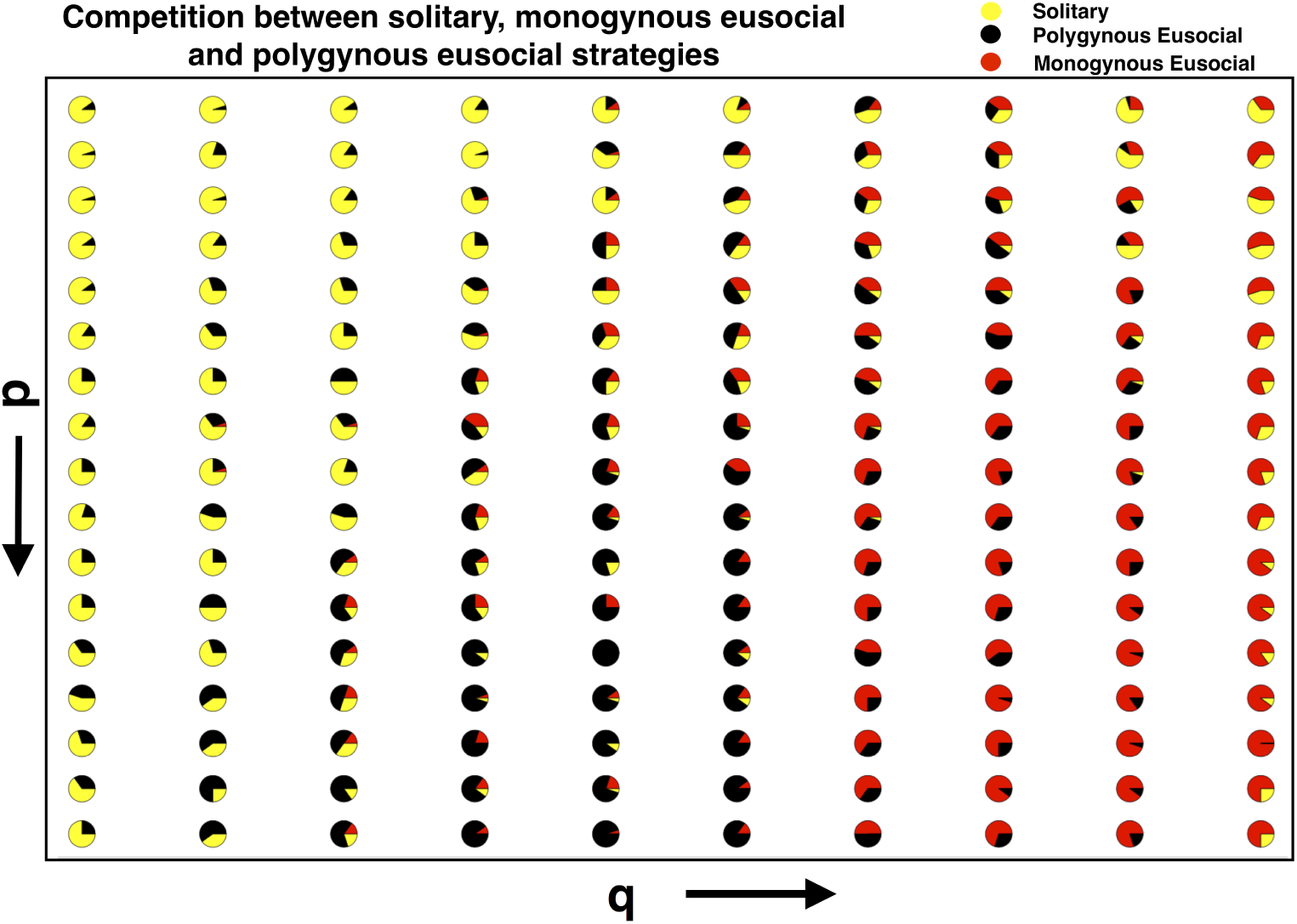
Competition between solitary, monogynous eusocial and polygynous eusocial populations: a unimodal reproduction profile is assumed, and the stay-put probability *q* is varied along the abscissa and the the maximum reproductive age *p* is along the ordinate. Monogynous eusociality, which evolves for larger values of *p* and *q* is depicted in red, the solitary strategy (in yellow) arises for lower values of *p* and *q*, while the black dots correspond to polygynous eusociality which comes about at intermediate values.

This would suggest that within this model, the evolution of eusociality and long lifespans are correlated. Short lifespans (low *p*) favour the solitary lifestyle, while for long *p* eusociality (whether polygynous or monogynous) is preferred. For values of *q* where all three strategies are possible, increasing *p* favours the monogynous strategy. This is in accord with the observation that polygynous species can have shorter lifespans compared to monogynous species with similar extrinsic death rates.

## Summary and Discussion

Classical theories of the evolution of ageing have limitations [16–18, 38–41]; empirical observations show that in some cases there are departures from classical predictions [16, 17], and mechanistic details of the evolution of ageing in many populations is generally unknown [18]. Furthermore, ageing is a process of considerable complexity [42–46]. Several earlier studies have quantitatively shown that the evolution of eusociality and that of long lifespans are highly correlated [8] although the mechanistic details need further exploration. Newer approaches such as the study of hierarchical trade-off models [20] and the inclusion of intergenerational transfer [47] are being employed to better understand such details. We therefore decided to explore the effect of a population structure on the evolution of eusociality.

In the present work, we have built upon an evolutionary model of eusociality that implements population structure [30] to include age structure. Using an age-structured NTW model and a related agent-based model, we show that the eusocial population structure can increase the fitness of long lifespan strategies. With fixed reproductive capacity in a lifetime and with extrinsic death in addition, strategies that favour slow senescence in solitary species are evolutionarily expensive. When there are considerable group benefits that accrue for longer lifespans, eusocial species outcompete solitary species. On the other hand, when the lifespan is short solitary species outcompete eusocial ones. The analysis of the fitness landscape for eusocial populations shows that with increased extrinsic death-rate, a population can increase fitness both by increasing as well as by reducing the intrinsic lifespan.

The comparison between fitness landscapes further shows that the evolution of long lifespans is possible over a larger region of parameter space for monogynous rather than polygynous eusocial populations. In polygynous eusocial species — intermediate between single queen colonies and solitary species — additional queens that join a group by migration effectively pull down the average age of colonies. The progeny can gain group benefits by joining another group which has already crossed a threshold size and so can increase early reproduction rate. In such cases, comparatively faster ageing (or shorter lifespans) would be beneficial. The ageing strategy also controls the number of colonies that exceed a threshold size for group benefits. Since the correlation is inverse, if the ageing rate of polygynous species is increased (namely a reduced lifespan) there can be a situation when there are no groups available which are above the threshold size. Progeny will then not be able to draw upon the benefits of joining a mature colony, and a solitary strategy would be preferred for very short lifespans. A similar argument can be made to show that for long lifespans the polygynous populations are dominant when the probability of progeny migration is large, and monogynous eusocial populations are dominant when this probability is small.

The competition simulations between long lifespan strategies show that the eusocial populations outcompete solitary population for a larger parameter space of probability to stay *q* and group benefits *g*. In our agent-based modelling simulations, the inclusion of polygyny along with the solitary and eusocial strategies allows for a better exploration of the parameter space for evolution of eusociality. Polygynous eusociality occupies an intermediary region in the phase space, between the solitary and monogynous eusocial cases. The evolution of long lifespans is thus intrinsic to eusociality, both evolving together. Indeed, it would appear from the present study that in this model a long lifespan in social organisms is both a product as well as an enabler of eusociality. This would suggest that including population structure in ageing models is important and further stresses the importance of heterogeneous modeling approaches.

In future work we intend to study extensions of the present models that can include more realistic forms of various age- and size-dependent parameters such as group benefits and death rates. A number of other features such as food availability, maturation time, or foraging time can be included, and their effects need to be explored since some of these factors are empirically known to affect ageing. Similarly, specific population and life-history structures such as the mating behaviour, maturation stage, inter-generational co-operation, metabolism and maintenance, need to be included and explored. the models themselves can be made more sophisticated by including processes such as mutation-selection.

Due to generality of the present models, specific quantitative predictions are difficult to make. However, the qualitative and theoretical results obtained here should ideally find validation through empirical observations. Nevertheless, eusociality is always correlated with long lifespans [8], and polygynous eusocial species have a lower lifespan compared to a monogynous one for a similar extrinsic death rate [10]. Understanding the mechanisms of ageing will have profound implications in medical interventions for senescence-related damage. In addition, insights into the ageing process will have an impact on our understanding of the dynamics of populations and their evolutionary trajectories.

## Acknowledgments

D. Raviteja thanks the CSIR, India for the Shyamaprasad Mukherjee Fellowship. Ram Ramaswamy holds the JC Bose National Fellowship of the Department of Science and Technology, India. We thank the referees for their critique and suggestions that have significantly improved this paper and have greatly helped us in our analysis.

